# Maximum tree lifespans derived from public-domain dendrochronological data

**DOI:** 10.1101/2022.08.13.503649

**Authors:** Franco Biondi, David Meko, Gianluca Piovesan

## Abstract

The public-domain International Tree-Ring Data Bank (ITRDB) is an under-utilized dataset to improve existing estimates of global tree longevity. Since dendrochronologists have usually targeted the oldest trees in a stand, this public-domain resource is bound to offer better estimates of maximum tree age than those available from randomized plots or grid-based forest inventories. We used the longest continuous ring-width series of existing ITRDB collections as an index of maximum tree age for that species and site. Using a total of 3679 collections, we obtained longevity estimates for 236 unique tree species, 156 conifers and 80 angiosperms, distributed all over the world. More than half of the species (167) were represented by no more than 10 collections, and a similar number of species (144) reached longevity greater than 300 years. Maximum tree ages exceeded 1000 years for several species (22), all of them conifers, while angiosperm longevity peaked around 500 years. As new collections are constantly being added to the ITRDB, estimates of tree longevity may change slightly, mainly by adding new species to the database. Given the current emphasis on identifying human-induced impacts on global systems, detailed analyses of ITRDB holdings provide one of the most reliable sources of information for tree longevity as an ecological trait.

**Key Message:** Baseline information on tree longevity was derived from the most extensive dendrochronological database currently available. The resulting summary provides a reference point, to be used for modeling and research purposes.

## Introduction

Tree longevity is an essential ecological trait for understanding forest vegetation dynamics (Gutsell and Johnson 2002), climatic impacts on woody species (Locosselli et al. 2020), and terrestrial carbon cycling (Körner 2017). While there is no research program specifically designed to investigate tree longevity, all research efforts aimed at predicting the fate of terrestrial ecosystems depend, more or less explicitly, on understanding and quantifying demographic patterns and traits, which include maximum tree lifespans. The emphasis currently being placed on modeling the future response of forest stands to climatic changes (especially atmospheric warming) and disturbance events (from droughts to wildfires) has prompted researchers to investigate resilience and resistance of woody species (e.g., Hessburg et al. 2019; Vitasse et al. 2019). In this context, as tree mortality is complementary to tree longevity (Das et al. 2016), baseline information on maximum tree age provides an index whose variability in time and space can reveal environmental and human impacts on forest species (Xu and Liu 2021).

For our purposes, tree age is defined as stem (or trunk) age, which is the cumulative duration of secondary growth since pith formation at a specified height from the ground (Piovesan and Biondi 2021). Using this definition, tree longevity can be determined for wood-forming species, either clonal or non-clonal, by means of dendrochronological methods, radiocarbon dating, or a combination of both (e.g., Piovesan et al. 2018). Dendrochronological work has been traditionally focused on the oldest individuals of a species, but existing tree-ring data has rarely been analyzed in terms of potential maximum lifespans. For instance, Zhao et al. (2019) reviewed in detail the International Tree-Ring Data Bank (ITRDB) and quantified its holdings in terms of species representation, spatial distribution, and potential improvements for macroecological research purposes, yet they did not address the issues connected with maximum tree lifespans.

Our objective was therefore to investigate tree longevity using the information contained in the holdings of the ITRDB, which are publicly available but are not yet searchable for demographic information. We present in this short communication the results of our analysis as a contribution to existing trait databases. An in-depth analysis of this information in terms of its significance for tree eco-physiology and evolutionary ecology is ongoing, and will be the subject of future publications.

## Materials and Methods

Ring-width data were obtained from the public-domain ITRDB repository in mid-March 2022. To enhance replication, we did not introduce any additional information besides what was available on the ITRDB ftp server (ftp.ncdc.noaa.gov/pub/data/paleo/treering). The four-letter codes that are traditionally based on the first two letters of the scientific (Latin) genus and species names (binomial nomenclature) were compared with their original meanings (Grissino-Mayer 1993).

The maximum length of all samples included in a collection was used to estimate tree longevity. To evaluate how reliable this index was, we compared it with a more refined estimate of longevity that was based on first grouping individual samples by tree code. This analysis was performed on a subset of the data, including 519 collections from Canada, Africa, and the Updates subdirectory. The maximum number of tree rings in a continuous sample exactly matched the tree-based estimate in most cases, with differences only in 64 collections, and with only two of them being greater than 100 years (**Figure S1**).

Additional checks were performed on the species name to avoid duplicates, incorrect entries, and collections where only the genus was given. A final comparison was made between the maximum sample length of a collection and the difference between the overall first and last year, which is included in the standard metadata information for each collection. When this difference exceeded the maximum series length by more than 100 years, we analyzed the collection using the COFECHA software (Grissino-Mayer 2001; Holmes 1983).

Summary statistics were calculated for all species as well as for angiosperms (Magnoliophyta) vs. conifers (Pinophyta), and also for the extra-tropics, defined as the regions with latitude above 30°N or below 30°S. Given that the tropics are in reality between 23.5°S and 23.5°N, other definitions could have been applied, such as ±25°, but we adopted Locosselli et al. (2020)’s definition for comparison purposes. In order to quantify the minimum number of sites that are most likely to generate reliable estimates of tree longevity, we tested the correlation between maximum tree age and number of ITRDB collections. All numerical analyses were performed using either the R numerical environment (R Core Team 2020) or the SAS software (Delwiche and Slaughter 2019).

## Results and Discussion

Overall a total of 3679 out of 5444 collections could be analyzed, which is a considerably larger number than previous summaries of ITRDB holdings (e.g., 2624 in Locosselli et al. 2020). The excluded files were affected either by non-standard data organization, end-of-line and end-of-record issues that could not be resolved, or both. The most recent ITRDB collection used to identify tree longevity was made in 2019, and the oldest one in 1978. Many species (76) were only represented by one collection, more than half of the species (167) were represented by no more than 10 collections, and a handful of species (7) were represented by more than 100 collections, with Douglas-fir (*Pseudotsuga menziesii*) being the species with the highest number (311).

Longevity estimates were obtained for 236 unique tree species, 156 conifers (3033 collections) and 80 angiosperms (646 collections), distributed all over the world but with greater density in the mid- and high-latitudes (**Figure 1** and **Appendix**). Areas with latitude above 30° N or below 30° S included 194 species, of which 65 were angiosperms (9 in the southern hemisphere) and 129 were conifers (19 in the southern hemisphere). The majority of species (144) reached longevity greater than 300 years, and maximum tree ages exceeded 1000 years for several species (22), all of them conifers, while angiosperm longevity peaked around 500 years (**Figure 2** and **Table 1**). This very large taxonomic difference in realized longevity is well known, albeit its causes are still being investigated (Munné-Bosch 2018; Peñuelas and Munné-Bosch 2010; Piovesan and Biondi 2021). Based on stochastic modeling of the theoretical relationship between average mortality rate and age structure in old-growth forests, maximum tree ages of a few centuries in angiosperms and of a few millennia in conifers are consistent with differences in their average mortality rate (Cannon et al. 2022).

**Table 1.** Summary of maximum tree ages estimated from ITRDB collections.

| Taxa | Species | Sites | Min-Max | Mean | St.Err.Mean | St.Dev. | Median | IQR<sup>a</sup> |
| --- | --- | --- | --- | --- | --- | --- | --- | --- |
|  | (#) | (#) | (yrs) | (yrs) | (yrs) | (yrs) | (yrs) | (yrs) |
| Angiosperms | 80 | 646 | 28-518 | 229 | 15 | 132 | 194 | 119-331 |
| Conifers | 156 | 3033 | 64-3205 | 618 | 40 | 496 | 504 | 313-770 |

<sup>a</sup>IQR: Inter-Quartile Range (1<sup>st</sup>-3<sup>rd</sup> quartile).

**Figure 1.**
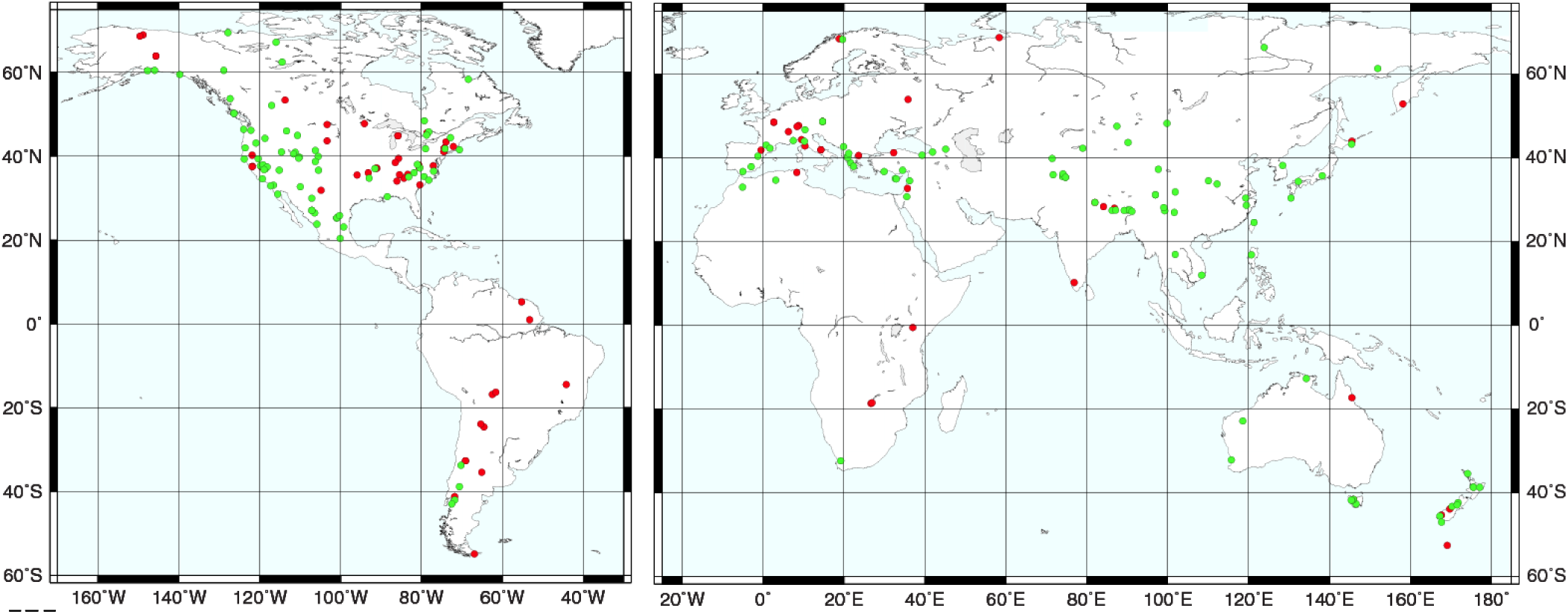
Map of 236 ITRDB collections (solid dots) that provided the maximum estimated tree age by species (80 angiosperms: red; 156 conifers: green). Sites cover most of the globe, from the Arctic (69.5 °N) to the sub-Antarctic (54.9 °S, Campbell Island), but with higher density in the extra-tropics (i.e., areas with latitude above 30° N or below 30° S), which included 194 species.

**Figure 2.**
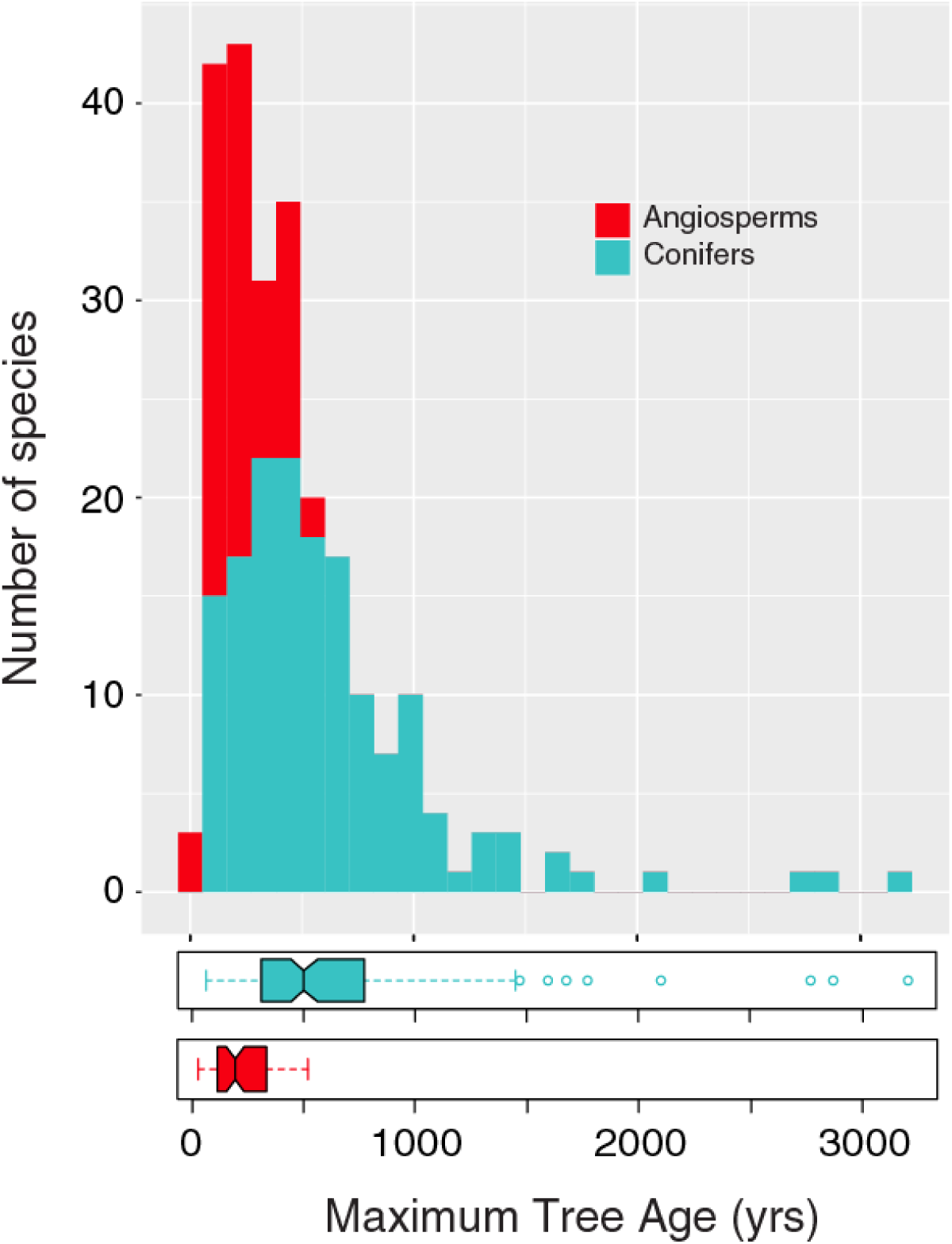
Distribution of tree longevity estimates, showing differences between angiosperms (red bars and boxplot) and conifers (green bars and boxplot).

Since dendrochronologists have usually targeted the oldest trees in a stand, the ITRDB public-domain data are bound to offer better estimates of maximum tree age than those available from randomized plots, grid-based inventories, or the most complex, state-of-the-art simulation models. As an example, based on a global analysis of forest inventories and climate data, Besnard et al. (2021) defined as “old growth” any stand older than 300 years, which is an order of magnitude less than the maximum tree ages we uncovered. While several large geographic regions were not included in Besnard et al.’s global analysis (“Africa, Indonesia and Australia were either underrepresented or not represented”), the authors recognized that even in regions where data were relatively abundant, such as the US, “unmanaged forests in remote areas were very likely less represented than managed forests”. The ITRDB data, as shown in our relatively simple analysis, therefore demonstrate that non-dendrochronological peer-reviewed approaches can severely underestimate tree longevity as an ecological trait.

We also note that there is an over-abundance of popular reports, either in press or on the internet, that exaggerate the age of the oldest trees, as it becomes clear when only dendrochronological or radiocarbon-based estimates are considered (Liu et al. 2022; Piovesan and Biondi 2021). Occasionally these unscientific claims are repeated in the most prestigious scientific journals, as shown by recent news that oaks older than a millennium can be found in the United Kingdom and in Fennoscandia (Pennisi 2022; Sonne et al. 2021). Denmark’s King Oak (*Quercus robur*) is an example of charismatic megaflora, but the notion that it could be “around 1,900 years of age” is nothing more than myth when confronted with science-based maximum reported ages of angiosperms in general, and of oaks in particular. Unrealistic tree ages, especially for very large stems, have often been obtained by assuming a constant growth rate (i.e., a constant ring width), calculated using only the outermost wood increments, which are typically smaller than the previous ones. Thus, Nunziata et al. (2022) could proclaim an estimated age of 2000-3000 years for the monumental chestnut (*Castanea sativa*) named “Castagno dei Cento Cavalli”, possibly favoring the local tourist industry, but not the scientific understanding of tree longevity.

Given that not all tree-ring data ever collected are deposited in the ITRDB, we performed an in-house evaluation of some species’ maximum tree ages using collections that we developed but have not yet been properly archived. Chronologies that have been published in connection with research projects in the Sierra Nevada (Meko et al. 2014) and the Great Basin (Biondi 2014) of North America provided estimates that in some cases exceeded the ITRDB ones, but ultimately did not result in large changes. For instance, single-needle pinyon (*Pinus monophylla*) reached 784 years (ITRDB: 653 yrs; see Appendix), big-cone Douglas-fir (*Pseudotsuga macrocarpa*) peaked at 683 years (ITRDB: 658 yrs), and blue oak (*Quercus douglasii*) topped at 539 years (ITRDB: 496 yrs). On the other hand, a large difference emerged for *Fagus sylvatica* (ITRDB: 407 yrs), which has a tree-ring-based maximum age of 625 yrs (Piovesan et al. 2019).

As new collections are constantly being added to the ITRDB, estimates of tree longevity may change. The comparison reported in the previous paragraph suggests that these changes may be relatively small for species that are already represented by several sites. Based on the relationship between estimated longevity and number of collections (**Figure 3**), ∼40 chronologies are needed to reach reliable estimates. Also, a larger percentage of angiosperm species appeared capable of reaching the extreme longevity of Magnoliophyta (a few centuries) compared to Pinophyta, since only a handful of species can attain the conifer maximum ages (a few millennia). Considering the very large number (3679) of ITRDB collections we analyzed, and that our results included 20% more species for the extra-tropics than previously reported (161; Locosselli et al. 2020), it is plausible that most changes in tree longevity estimates derived from tree-ring data will be caused by adding new species to the ITRDB holdings. Yet, we note that our overall estimated mean longevity of trees in all extra-tropical biomes was 516 ± 34 yrs, which is significantly higher than the recently published estimate of 322 ± 200 yrs (Locosselli et al. 2020).

**Figure 3.**
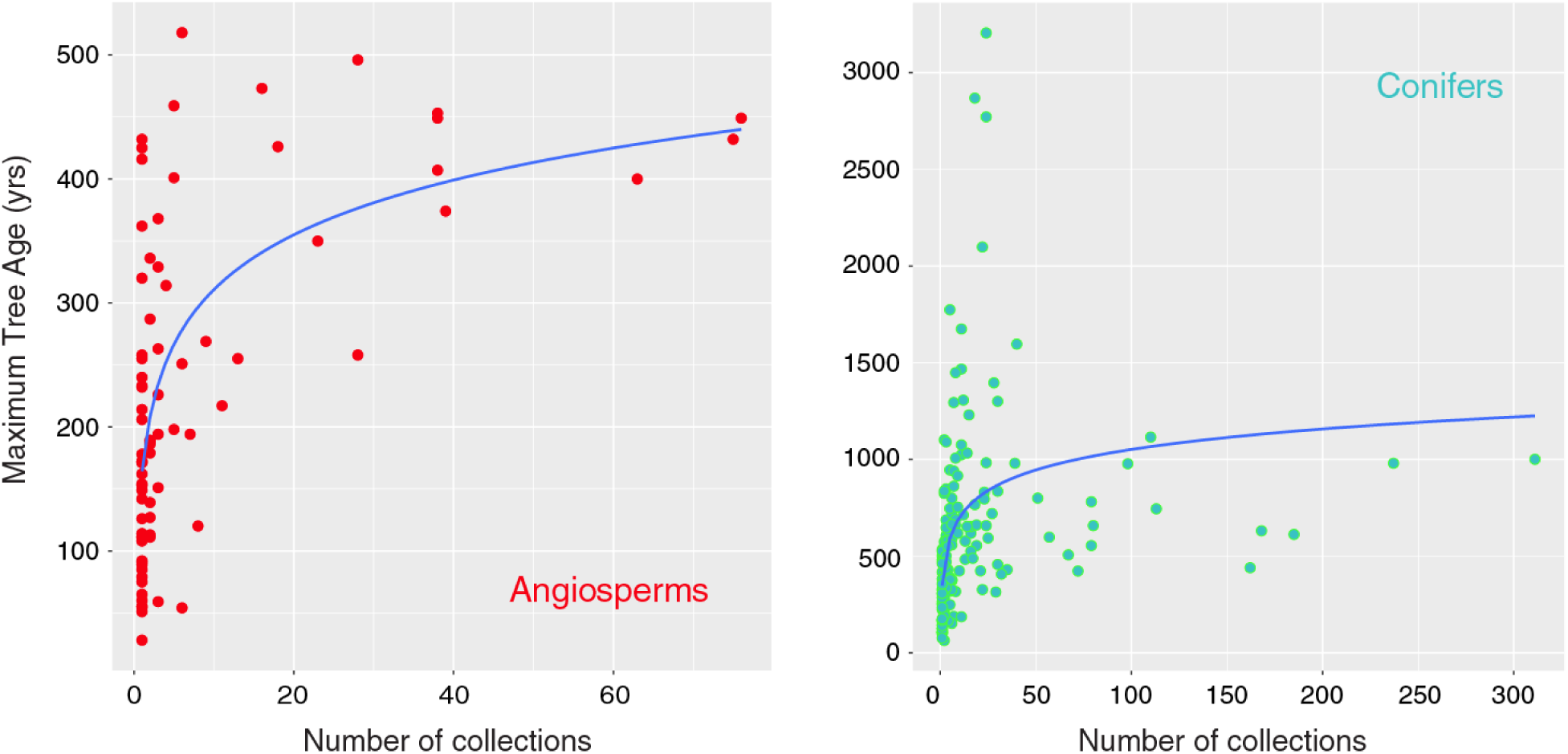
Relationship between the estimated tree longevity (Maximum Tree Age) and the number of collections for each species (80 angiosperms, 156 conifers).

Despite the advantages of tree-ring records for estimating tree longevity, it is still necessary to point out that the scientifically-based data we have produced on such a fundamental botanical and ecological trait represent the minimum boundary for a species. In some cases, tree-ring samples may contain many more rings that are not measured, and are therefore excluded from ITRDB holdings. Furthermore, dendrochronologists may often avoid measuring sections of increment cores or stem sections that are too difficult to crossdate, either because of erratic growth patterns, extremely low growth rates, injuries, branch insertions, rot, or other anatomical imperfections of the wood structure (Piovesan and Biondi 2021).

An additional confounding factor is that, even when tree-ring measurements are archived in the ITRDB, researchers may not provide the entire datasets. By doing so, investigators can satisfy funding agency requirements for archiving data while at the same time avoiding to share the most important, i.e. longest-term, information. This issue was noticed in more than one case, but a clear example was provided by the 37 California chronologies coded as CA561-CA597, which all end in 1990-1991 and start in 1879-1880. Since the collections only cover 111-112 years, but were made on species (*Abies concolor, A. magnifica, Calocedrus decurrens, Pinus contorta, P. jeffreyi, P. lambertiana, P. ponderosa, Tsuga mertensiana*) and in areas (the Sierra Nevada of the western USA) that are known to yield much older trees (see Appendix), it is unlikely that all data were archived. One could argue that perhaps the study was performed in even-aged plantations, or that there were special constraints that forced the investigators to sample young trees or to extract very short increment cores even when the stem was large. As it turns out, one of us (FB) actually participated in some of those field collections as a graduate student, acquiring first-hand knowledge of these stands and of these collections, which were dendroclimatic-oriented and performed in old-growth stands by targeting the largest trees.

When the number of ITRDB collections of the same species is large enough, the above mentioned issue should not impact the estimated maximum tree age. However, a potentially large underestimation occurs if data are not fully archived and only a few chronologies are available for a species. Among the collections coded as CA561-CA597 are indeed the only ITRDB holdings for a species, *Quercus kelloggii*, whose longevity was therefore estimated at 111 years – an unreliably small value. Partial submissions may cause other artifacts, for instance connected to changes in tree longevity over time. While we did not perform an exhaustive analysis of this problem, one can imagine how the maximum age of tree species included in collections CA561-CA597 could be compared to the longevity of the same species in earlier collections. As reports of the impending doom of ancient trees accumulate (Locosselli et al. 2020; McDowell et al. 2020), such a comparison could then lead to claims of human-induced reduction in tree longevity even without the presence of a naive observer or one fully vested in promoting an apocalyptic narrative.

The definition of ‘old-growth’ stands, which has fundamental implications for conservation efforts and science-based forest management, depends on correctly estimating tree longevity. We emphasize that what ‘old’ means depends both on the tree species, as shown here, and on its realized niche, as we have argued elsewhere (Piovesan and Biondi 2021). Using a fixed cutoff, such as the 300 years threshold that is often repeated in the literature (e.g., Besnard et al. 2021), fails to consider ecoclimatic and taxonomic differences. Earlier, detailed analyses of old-growth conditions had already pointed out that reported old-growth forest ages can range from 50 to 1,150 years (Wirth et al. 2009), making it necessary to design new metrics for evaluating old-growth conditions (Di Filippo et al. 2017). Additional submissions of tree-ring data to the ITRDB, and related publications of dendrochronological and radiocarbon-based information on tree longevity, is bound to improve our understanding of tree life histories, forest demographics, old-growth features, and of their complex dependence on multi-scale impacts from natural and human-caused disturbances.

## Acknowledgments

We are grateful to the Contributors of the International Tree-Ring Data Bank, and to the agencies and institutions that have allowed the establishment and maintenance of this exceptional, publicly available resource.

## Statements & Declarations

### Author Contributions

All authors contributed to the study conception and design. Data collection and analysis were performed by F. Biondi with contributions by D. Meko and G. Piovesan. The first draft of the manuscript was written by F. Biondi, and all authors commented on previous versions of the manuscript. All authors read and approved the final manuscript.

### Data Availability

The datasets generated during the current study are available from the corresponding author on reasonable request.

**Figure S1.**
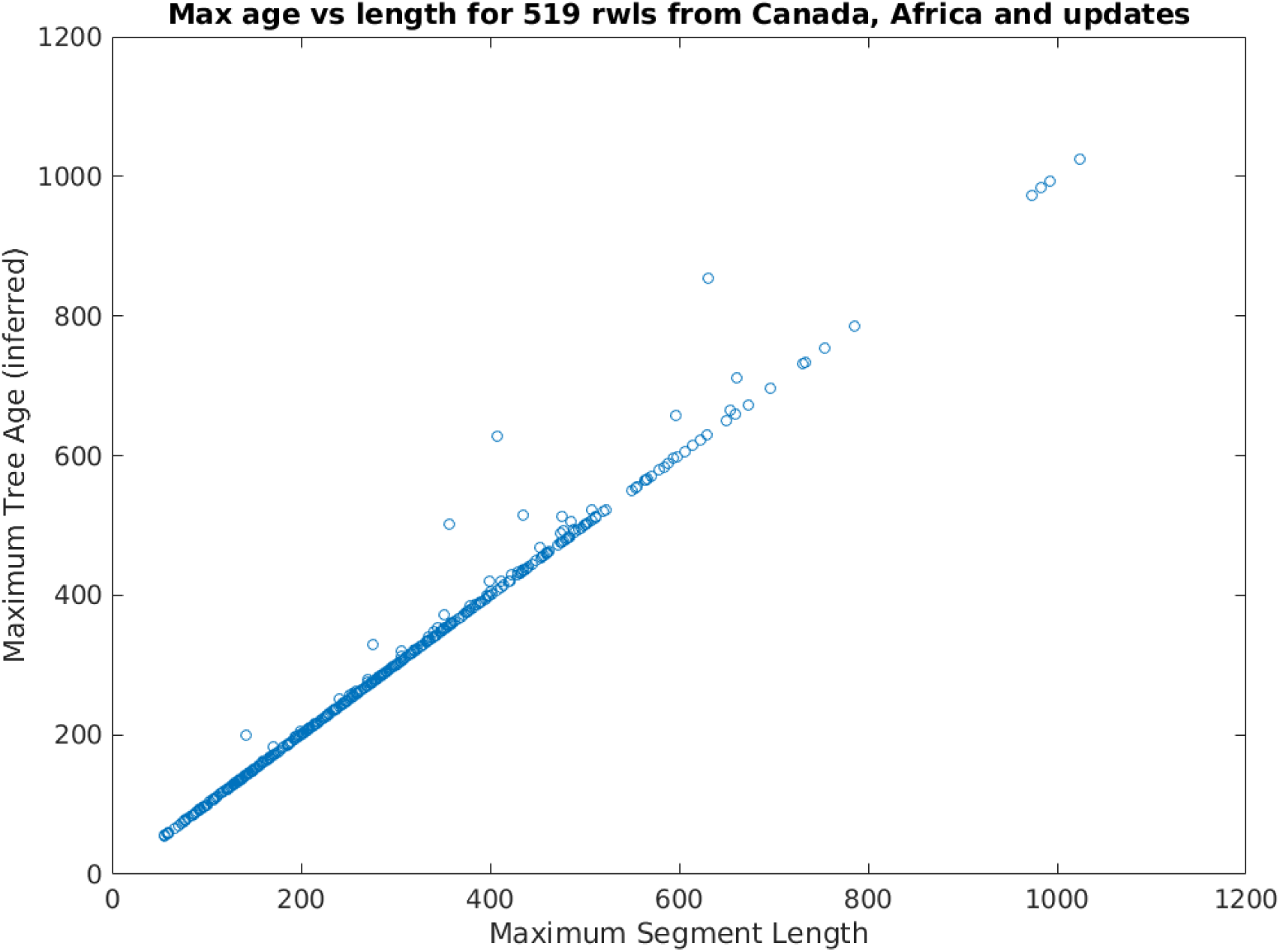
Comparison between tree ages estimated using the maximum length of a single continuous ring-width measurement series (x-axis) and the maximum length of ring-width measurements for a tree (y-axis), as coded in 519 ITRDB collections from Canada, Africa, and the Updates subdirectory in March 2022.

